# Culture system for longitudinal monitoring of bone remodeling *ex vivo*

**DOI:** 10.1101/2023.10.12.562045

**Authors:** E.E.A. Cramer, K.C.J. Hermsen, L.M. Kock, K. Ito, S. Hofmann

**Affiliations:** Orthopaedic Biomechanics, Department of Biomedical Engineering and Institute of Complex Molecular Systems, Eindhoven University of Technology, P.O. Box 513, 5600 MB Eindhoven, the Netherlands; Institute of Complex Molecular Systems (ICMS), Eindhoven University of Technology, P.O. Box 513, 5600 MB Eindhoven, the Netherlands; LifeTec Group BV, Eindhoven, The Netherlands

## Abstract

To quantify and visualize both bone formation and resorption within osteochondral explants cultured *ex vivo* is challenging with the current analysis techniques. An approach that enables monitoring of bone remodeling dynamics is longitudinal micro-computed tomography (µCT), a non-destructive technique that relies on repeated µCT scanning and subsequent registration of consecutive scans. In this study, a two-compartment culture system suitable for osteochondral explants that allowed for µCT scanning during *ex vivo* culture was established. Explants were scanned repeatedly in a fixed orientation, which allowed assessment of bone remodeling due to adequate image registration. Using this method, bone formation was found to be restricted to the outer surfaces when cultured statically. To demonstrate that the culture system could capture differences in bone remodeling, explants were cultured statically and under dynamic compression as loading promotes osteogenesis. No quantitative differences between static and dynamic culture were revealed. Still, only in dynamic conditions, bone formation was visualized on trabecular surfaces located within the inner cores, suggesting enhanced bone formation towards the center of the explants upon mechanical loading. Taken together, the *ex vivo* culture system in combination with longitudinal µCT scanning and subsequent registration of images demonstrated potential for evaluating bone remodeling within explants.

## 1. Introduction

Bone is a dynamic tissue that continuously remodels its structure in response to mechanical stimuli present in its environment. This mechanotransduction process is governed by the highly regulated interplay between osteocytes, osteoblasts, and osteoclasts. Osteocytes, which are embedded within the mineralized matrix, detect changes in their surroundings, which they transmit to other cells, including osteoclasts and osteoblasts (Clarke, 2008; Robling et al., 2006). In this way bone remodeling, which involves the removal of mineralized bone by osteoclasts followed by the formation of bone matrix through the osteoblasts, is coordinated. The culture of bone fragments explanted from living bone provides a unique method to study remodeling processes with bone cells maintained within their native ECM (Marino et al., 2016). Within bone explants, osteocytes remained in their lacunar spaces (Dallas & Moore, 2020) and have been shown to preserve viability and functionality for a period of at least 14 days (Chan et al., 2009; Davidson et al., 2012). In addition, the application of mechanical loading onto bone explants revealed osteocyte responses such as PGE_2_, PGI_2_ and Ca^2+^ release known as early events in the cascade of mechanotransduction (Chan et al., 2009; Haj et al., 1990; Ishihara et al., 2013; Jing et al., 2014; Morrell et al., 2020; Rawlinson et al., 1991, 1995). The release of these biochemical factors was found to be related to induced bone formation after 4 weeks (Birmingham et al., 2015a, 2016; Chan et al., 2009; Mann et al., 2006). Despite the unique environment in explants to study bone turnover, the measurement and visualization of bone remodeling processes ex vivo, preferably over time, remains challenging with the current analysis techniques.

Current techniques that are used to evaluate remodeling in bone explant cultures include the measurement of bone markers in culture supernatant and histological evaluation (Davies et al., 2006; Klüter et al., 2020; Schnieders et al., 2013; Stoddart et al., 2006). Examining biomarkers can provide insight into overall bone turnover, but it does not provide specific information about local processes taking place. In contrast, histology enables visualization of local processes at a specific stage but requires many samples to be sacrificed if multiple timepoints need to be investigated. Dynamic histomorphometry is another analysis technique applied in *ex vivo* cultures and allows for both visualization and quantification of bone formation over time (Birmingham et al., 2016; Curtis et al., 2018; David et al., 2008; Mann et al., 2006). The administration of calcium binding fluorochromes at specific timepoints during culture reveals newly mineralized bone matrix, enabling quantification of bone formation by utilizing distances between fluorochrome labels and the time interval between their administration. A disadvantage of this method is that it is a semi-quantitative analysis because mineralization rates are obtained from images taken at specific locations and do not represent the complete sample. Moreover, samples need to be destroyed before evaluation and it is not possible to examine bone resorption, which is the other key component of bone remodeling. The drawbacks of the current methods highlight the need for implementation of different techniques to measure bone remodeling in bone explant cultures.

A non-destructive method to capture and quantify both bone formation and resorption is the repeated application of micro-computed tomography (µCT) (van Lenthe et al., 2007; Wehrle et al., 2019). Registration of consecutive µCT images allows for three-dimensional assessment of bone turnover (Christen & Müller, 2017; Wehrle et al., 2019). Specifically, voxels appearing only in the first scan are considered resorbed bone and voxels present only in the follow-up scan are considered formed bone. *In vivo*, this method has demonstrated to be successful in studying remodeling and fracture healing in both animals and humans (Boyd et al., 2006; Christen & Müller, 2017; David et al., 2003; Lambers et al., 2012; Schulte et al., 2011). Regarding *ex vivo* cultures, one study by Birmingham et al., utilized this technique of registration of µCT images taken pre- and post-experiment and showed increased bone formation in trabecular bone explants exposed to vibrational loading compared to static controls (Birmingham et al., 2015a). Although bone formation and resorption were captured, limitations were reported with misalignment of registered images and motion artifacts during scanning because samples were scanned in different orientations for each scan. This resulted in registration errors and restriction of the volume that could be examined.

Most bone research performed *ex vivo* utilizes explants comprised of solely bone tissue where the cartilage layer was removed before culture, thereby ignoring the highly interconnected relationship between the two tissues. Explant cultures comprising both tissues, so called osteochondral explants, provide a closer representation of the *in vivo* joint-like environment as there is direct communication between the tissues. Thus far, osteochondral explants are mainly utilized as models to investigate cartilage repair treatment and to study diseases such as osteoarthritis, in which both tissues are affected partly (de Vries-van Melle et al., 2012, 2014; Duchi et al., 2019; Geurts et al., 2018; Kleuskens et al., 2021; Maglio et al., 2019; Schwab et al., 2017). Schwab et al. addressed the issue of having two tissues requiring different culture conditions. They developed a culture platform comprising two compartments, one for bone and one for cartilage, in which each tissue received its respective medium (Schwab et al., 2017). Using their platform, it was shown that the bone part could remain viable over a culture period of 12 weeks. With their aim to study cartilage treatment strategies, further evaluation on processes occurring in the bone part, such as spatiotemporal analysis of bone dynamics, was not performed. Moreover, the two-compartment culture system established by Schwab et al. does not allow for evaluation of bone remodeling over time using µCT because osteochondral explants should be taken out of the culture plate each time to perform a µCT scan. This could lead to different orientations in each scan and the transfer of the samples increases the risk of contaminations.

Gaining insight into bone remodeling processes could enable osteochondral cultures to be utilized as model to study novel therapies that affect formation and/or resorption processes and diseases affecting both tissues. Therefore, this study aims to establish a two-compartment culture system for osteochondral explants that is compatible with µCT scanning thereby enabling longitudinal monitoring of bone formation and resorption during culture. Without the need to take samples out of the culture system before each scan, it is expected that ensuring a similar position during each scan will reduce the risk of errors and misalignment during registration, thereby enabling registration of the complete volume of the explant. To demonstrate that the culture system could capture differences in bone remodeling, in the second part of this study we aimed to implement compressive mechanical loading to enhance bone formation *ex vivo* (Birmingham et al., 2016; David et al., 2008; Lozupone et al., 1996; Mann et al., 2006). To ensure imposed compressive forces are received by the bone tissue, bone explants without a cartilage layer were utilized, because cartilage will alter loading patterns experienced by the bone. This makes culture in a two-compartment system unnecessary. Hence, our culture system was designed to be adjustable for scanning of bone explants also in one compartment and thereby allows for monitoring of bone remodeling in both osteochondral and bone explants.

## 2. Materials & Methods

### 2.1 Isolation of porcine explants

Osteochondral explants were isolated from porcine femoral condyles, obtained freshly from the slaughterhouse’s left-over material. After removal of soft tissue and opening of an intact knee joint under aseptic conditions, femoral condyles were exposed. Using a hollow-drill (MF Dental, Weiherhammer, Germany) and a custom-made hollow metal tube serving as break off tool (**Figure S1A, C-D**) osteochondral cores with Ø 10 mm were isolated from the condyles. The drilling process was performed under constant cooling with sterile PBS (P4417, Sigma-Aldrich) containing 2% v/v antibiotic-antimycotic (anti-anti) (15240, Gibco^TM^, Thermo Fisher Scientific) at 4°C. The bone end was cut planar and orthogonal to the longitudinal axis using a bench saw (KS230, Proxxon, Föhren, Germany) to obtain a bone height of approximately 10 mm (**Figure S1E**). Osteochondral explants were obtained from 3 donors. Between steps, explants were stored in sterile PBS with 2% v/v anti-anti.

For the culture involving compressive mechanical loading, the isolation steps were performed as described above with small adaptations because explants that consisted only of bone tissue, without cartilage, were used. The explant size was adapted to fit into the compression bioreactor. Bone cores with Ø 8 mm were isolated. The cartilage layer was removed using a bench saw (KS230, Proxxon, Föhren, Germany). Subsequently, the bone end was cut using the bench saw in combination with a custom-made tool (**Figure S1B**) to obtain bone explants with a similar height of 8 mm. Bone explants were obtained from 10 donors.

### 2.2 Static culture of osteochondral explants

After harvesting, each osteochondral explant was mounted in an insert using an O-ring (Brammer, Leeds, UK) and placed in a culture chamber creating two separated compartments, with cartilage exposed to medium in the lower compartment and the bone part in the upper compartment (**Figure 1A-B**). The system was closed with a screw thread lid to ensure a sterile environment. This culture system made of polysulfone was adapted from the previous described two-chamber culture system for osteochondral explants in wells-plates (Schwab et al., 2017) and a culture system compatible with µCT scanning *in vitro* (Hagenmüller et al., 2007). The bottom chamber was filled with 5 ml of cartilage medium consisting of DMEM (hg-DMEM, 14966, Gibco^TM^, Thermo Fisher Scientific), 1% v/v anti-anti (15240, Gibco^TM^, Thermo Fisher Scientific), 40 µg/ml L-proline (Sigma-Aldrich), 50 µg/ml ascorbic-acid-2-phosphate (A8960, Sigma-Aldrich) and 1% v/v ITS+ premix (Corning, Thermo Fisher Scientific). The upper chamber contained 3.5 ml of bone medium consisting of DMEM (hg-DMEM, 14966, Gibco^TM^, Thermo Fisher Scientific), 10% v/v FBS (BCBV7611, Sigma-Aldrich), 1% v/v anti-anti (15240, Gibco^TM^, Thermo Fisher Scientific), 10 mM ß-glycerophosphate (G9422, Sigma-Aldrich), 50 µg/ml ascorbic-acid-2-phosphate (A8960, Sigma-Aldrich) and 100 nM dexamethasone (D4902, Sigma-Aldrich). Explants (n=5) were cultured for 6 weeks in an incubator at 37°C with 5% CO_2_ and medium was replaced 3 times a week in both compartments. Medium supernatant of the bone compartment was collected and stored at -80°C until further analysis.

**Figure 1.**
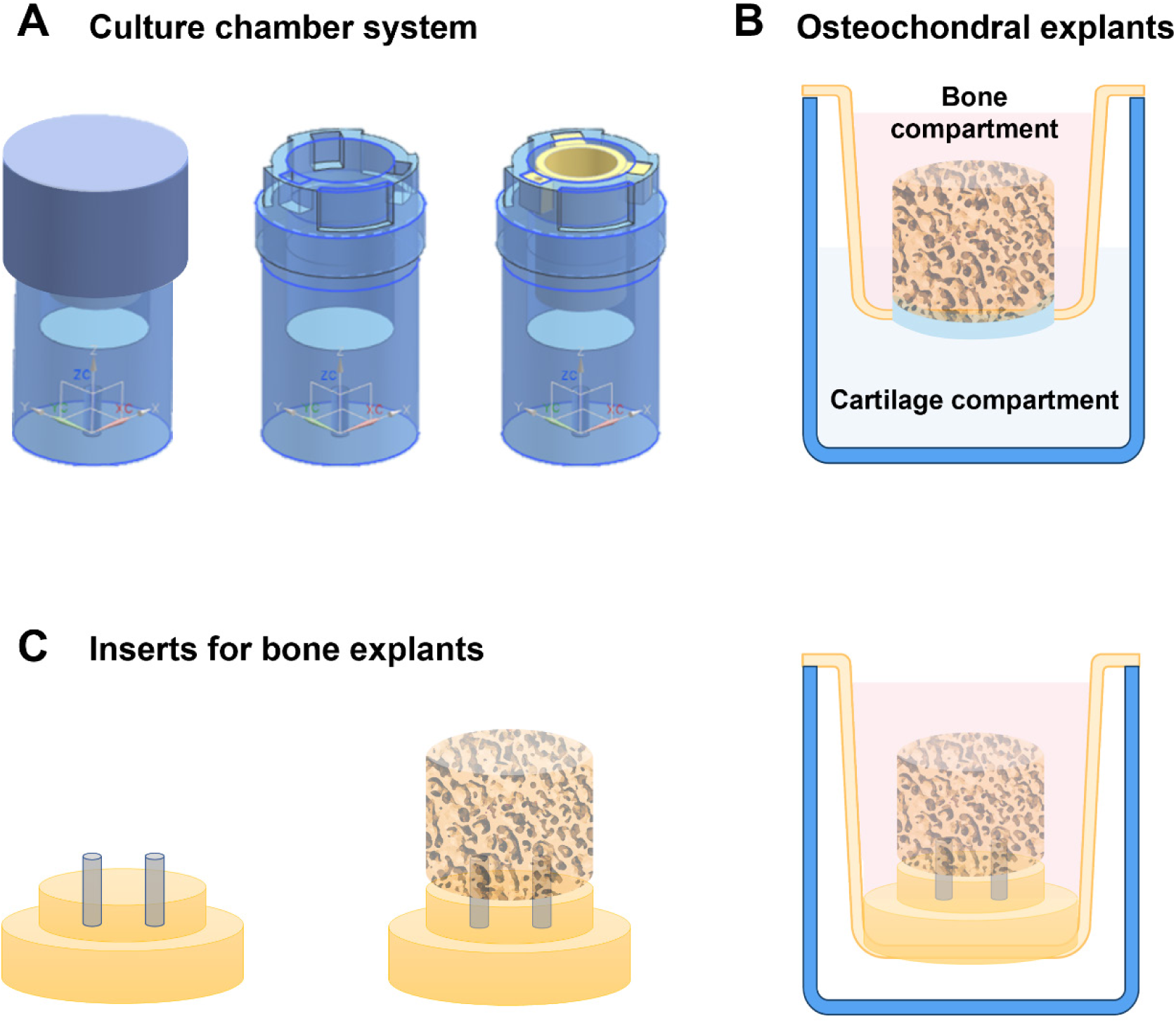
– Schematic of culture system compatible with µCT scanning. Culture chamber with insert (yellow) to create a two-compartment system to allow culture of osteochondral explants (A). The upper compartment contains medium specific for the bone part of the osteochondral explant and the lower compartment accommodates cartilage tissue with its respective medium (B). The culture system is adjustable for µCT scanning (and culture) of bone explants (without cartilage) when the second, lower compartment is redundant. Bone explants can be mounted on a holder with two pins which is clicked into the insert within the culture chamber (C). This ensures a fixed position during scanning and creates only one compartment that needs to be filled with bone medium. Osteochondral explants are 10 mm in diameter, bone explants are 8 mm in diameter. The complete culture system (blue) has a diameter of 50 mm.

### 2.3 Dynamic culture of bone explants

A compression bioreactor, developed by LifeTec Group BV (Eindhoven, the Netherlands), was used to apply dynamic compressive loading (**Figure S2**). The bioreactor system consisted of a linear actuator connected to a custom-made 6-well plate (**Figure S2A-C**). Bone explants were mounted in Ø 8 mm custom-made inserts (**Figure S2D**) to keep samples in place inside the well. Wells were sealed off with flexible lids where flat-ended stainless-steel pistons (Ø 10 mm) penetrated through (**Figure S2E**). The complete well-plate was covered with an extra lid (**Figure S2C**). The pistons allowed for adjustment of height per individual sample, which was adjusted before each loading procedure. Four explants were loaded at the same time, because the load is distributed over the loaded samples. Bone explants were subjected to 40 N of compression at a frequency of 1 Hz for 30 minutes per day, 5 days a week. This dynamic loading regime was chosen based on previous explant cultures (David et al., 2008) and the force value was based on estimations made using a Young’s modulus found in literature (∼300 MPa) for comparable bone cores (Li & Aspden, 1997) and a strain value in the window of bone formation according to the mechanostat theory (∼3000 microstrain) (Frost, 2003; Kan et al., 2014). A 5 N pre-load was applied. Displacement of the pistons during each loading regime and corresponding force were recorded by the bioreactor software and extracted in force-displacement curves.

Bone explants (n=8) were cultured in medium consisting of α-MEM (41061, Gibco^TM^, Thermo Fisher Scientific), 10% v/v FBS (Hyclone^TM^, GE Healthcare, USA), 1% v/v anti-anti (15240, Gibco^TM^, Thermo Fisher Scientific), 10 mM ß-glycerophosphate (G9422, Sigma-Aldrich) and 50 µg/ml ascorbic-acid-2-phosphate (A8960, Sigma-Aldrich) for 4 weeks at 37°C with 5% CO_2_. Medium was changed 3 times a week and collected on day 2, 4, 8, 10, 15, 17, 22, 24 and stored at -80°C until further analyses. In parallel, similar sized bone explants (n=8) were cultured in static condition in a regular well-plate for direct comparison between static and dynamic conditions.

### 2.4 Biomechanical testing

To determine the Young’s modulus of porcine bone explants (Ø 8 mm, height 8 mm), compression tests were performed using a tensile tester (Model 42, MTS Criterion, Eden Prairie, USA) equipped with a loadcell of 1 kN (LSB.103, MTS systems corp., Eden Prairie, USA). After a preload of 5 N was reached, samples were compressed at a strain rate of 1%/sec until a maximum force of 200 N. **Formula 1** was used to determine the compression modulus E [Pa], which was derived from the linear part of stress-strain curves. Stress σ [Pa] was derived from the applied force F [N] and the cross-sectional area A [m^2^] of the explant surface. Strain ε [-] was calculated from change in length ΔL [m] divided by the original length L [m].

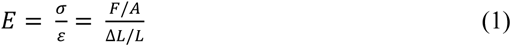

### 2.5 Measurement and visualization of bone formation and resorption

Osteochondral explants (n=5) were scanned weekly using micro-computed tomography (µCT) (µCT100, Scanco Medical, Brüttisellen, Switzerland). Culture chambers were specifically designed to allow µCT scanning during culture without the need to take samples out of their sterile culture environment and to ensure a similar orientation of the sample each time (**Figure 1A**). Scanning was performed at an energy level of 55 kVp, intensity of 200 µA and integration time of 200 ms, AL filter of 0.5 mm and a voxel size of 16.4 µm. For longitudinal image analysis, follow-up images were registered onto images of previous scans using an automated algorithm in the IPLFE software (Scanco Medical). Voxels appearing only in the first scan were considered resorption and voxels present only in the follow-up scan were considered formation. Bone formation (or gain) and resorption (or loss) were determined by finding clusters of ≥ 5 voxels that had a difference in attenuation coefficient between both scans ≥ 225 mgHA/cm^3^. These thresholds were chosen based on literature (Christen et al., 2014, 2018) and based on registration of one sample that was scanned repeatedly on the same day to examine noise levels. With these thresholds applied, noise levels below 1% of the total volume were obtained. Bone explants were segmented at a threshold of 450 mgHA/cm^3^ and a Gaussian filter was applied to all scans using a filter support of 1 and filter width sigma of 0.8. Bone formation (orange) and resorption (purple) were visualized with color-coded images and quantified as percentage of unchanged bone volume (grey) which appeared in both scans. Percentages of bone resorption were subtracted from bone formation to calculate net bone remodeling.

Bone explants used for the dynamic culture experiment (n=4 per group) were scanned on day 1 and day 28 using an adapted version of the culture chamber system (**Figure 1C**) with only one chamber. Using a guiding tool (**Figure S1A**), two holes (Ø 1 mm) were drilled, which allowed the explant to be placed on two pins, made from polyether ether ketone (PEEK) material, on a custom-made holder ensuring similar orientation with each scan. This holder was clicked into an insert which allowed the explant to maintain a fixed position during scanning. The insert, filled with culture medium, was placed in the larger culture chamber similar to the culture chamber for the osteochondral explants.

### 2.6 Visualization of tissue and cellular components in osteochondral explants

To examine structural components including cell nuclei and ECM within osteochondral explants, hematoxylin and eosin (H&E) staining was performed. At the end of the culture period, one half of the osteochondral explant (n=5) was fixed overnight in 10% neutral buffered formalin, washed in PBS and immersed in decalcification solution consisting of 10% w/v EDTA (E4884, Sigma-Aldrich) in water at pH 7. This solution was changed 3 times a week for a period of 4 weeks. After decalcification, samples were washed with water to remove remaining EDTA, and subsequently dehydrated through graded ethanol series and xylene in an automatic tissue processor (Microm STP-120, Thermo Fisher Scientific) and embedded in paraffin (P3558, Sigma-Aldrich). Sections of 6 µm thickness were sliced using a microtome (Leica RM2255, Germany), attached to poly-L-Lysine coated microscope slides (Thermo Fisher Scientific) and dried overnight. Slides were deparaffinized in xylene followed by decreasing series of ethanol (3x 100%, 96% and 70%), hydrated and stained with Mayer’s hematoxylin (MHS16, Sigma-Aldrich) and Eosin Y (HT110216, Sigma-Aldrich) according to the manufacturer’s instructions. After dehydration in graded series of ethanol (70%, 96%, 3x 100%) and xylene, stained sections were mounted with Entellan^TM^ (107961, Sigma-Aldrich) and imaged using a bright field microscope (Zeiss Axio Observer Z1, Germany).

### 2.7 Visualization of matrix formation in osteochondral and bone explants

To visualize newly formed matrix, RGB trichrome stain was performed on decalcified sections. RGB trichrome stain allows to discriminate between calcified and uncalcified bone and cartilage in sections of decalcified paraffin embedded tissues using sequential staining of three dyes, Picrosirius Red, Fast Green, and Alcian Blue (Gaytan et al., 2020). After culture, explants were cut in half, fixed, decalcified, dehydrated, embedded in paraffin, sectioned and deparaffinized as described above for H&E staining. Sections were stained in 1% w/v Alcian Blue (A5268, Sigma-Aldrich) in 3% v/v aqueous acetic acid (A6283, Sigma-Aldrich) solution for 5 minutes, followed by 1% w/v Fast Green (F7258, Sigma-Aldrich) in distilled water for 20 minutes and subsequently stained in 0.1% w/v Sirius Red (365548, Sigma-Aldrich) in picric acid solution (36011, Sigma-Aldrich) for 10 minutes. Between steps, sections were rinsed in tap water. After dehydration in graded series of ethanol (70%, 96%, 3x 100%) and xylene, stained sections were mounted with Entellan^TM^ (107961, Sigma-Aldrich) and imaged using a bright field microscope (Zeiss Axio Observer Z1, Germany). Day 0 osteochondral samples were stained with RGB dyes as reference for the start of culture.

### 2.8 Immunofluorescent visualization of mesenchymal stromal cells and osteoblasts

To evaluate the presence of mesenchymal stromal cells (MSCs) and osteoblasts, sections were stained for CD271, as marker for bone marrow MSC-like cells, and osteocalcin, as marker expressed by osteoblast-like cells. Prepared paraffin sections of osteochondral tissue were deparaffinized as described above and stained. Briefly, sections were immersed in 1x citrate buffer (S1699, Dako) overnight at 60°C for antigen retrieval. After a washing step in 0.05% v/v PBS-Tween (Tween 20, 822184, Merck), samples were blocked in 5% v/v normal goat serum in PBS for 1 hour. Primary antibodies for CD271 (1:100, 14-9400-82, eBioscience^TM^, Thermo Fisher Scientific) and osteocalcin (1:200, ab13418, Abcam) in 1% v/v normal goat serum in PBS were added to samples and incubated overnight at 4°C. Secondary antibody (1:200, A21240, Alexa 647, Thermo Fisher Scientific) in PBS was incubated on samples for 1 hour at RT followed by DAPI staining, 0.1 µg/ml in PBS, for 10 min. Coverslips were mounted with Mowiol and images were acquired with a laser scanning microscope (Leica TCS SP8X).

### 2.9 Measurement of cell metabolic activity

To determine cell metabolic activity, AlamarBlue^TM^ (A13262, Thermo Fisher Scientific) was added directly to the culture medium of bone explants in dynamic (n=4) and static condition (n=3) at a concentration of 10% v/v and incubated for 4.5 hours at 37°C, 5% CO_2_. After incubation, 100 µl of the medium was pipetted in a black 96-wells assay plate and fluorescence was measured with a plate reader (Synergy^TM^ HTX, Biotek, Winooski, VT, USA) at excitation 530/25 nm, emission 590/35 nm. Measured fluorescent values were corrected for fluorescent values of medium samples cultured without explants (blank). After the assay, samples were washed two times with sterile PBS at 37°C before fresh medium was added. The AlamarBlue^TM^ assay was performed on day 6, 13, 20 and 27.

### 2.10 Cell death measurement

Lactate dehydrogenase (LDH) activity, an indicator for cell death, was quantified in the culture supernatant taken 3 times a week during media change from the dynamically (n=6) and statically (n=6) cultured bone explants. Medium samples from day 2 and day 4 had to be diluted 1:1 in PBS because of the higher levels of cell death in the first days after harvesting, caused by the isolation procedure. 100 µl of medium supernatant was mixed with an equal volume of LDH reaction mixture, which was prepared according to the manufacturer’s instructions (11644793001, Roche, Sigma-Aldrich). Absorbance was measured directly after mixture and then every 10 minutes to ensure values were within standard curve range using a plate reader (Synergy^TM^ HTX, Biotek, Winooski, VT, USA) at 492 nm. Standards with known NADH concentrations (10107735001, Roche, Sigma-Aldrich) ranging between 0 and 0.75 µmol/ml were used to calculate the LDH activity per explant.

### 2.11 Measurement of tartrate resistant acid phosphatase activity

To examine osteoclast activity, tartrate resistant acid phosphatase (TRAP) activity levels were measured. TRAP activity was quantified in the culture medium of osteochondral explants (n=5) and bone explants in dynamic (n=6) and static (n=6) condition. P-nitrophenyl phosphate buffer was prepared by mixing 1 mg/ml p-nitrophenyl phosphate disodium hexahydrate (71768, Sigma-Aldrich) and 30 µl/ml tartrate solution (3873, Sigma-Aldrich) with stock buffer consisting of 0.1 M sodium acetate (S7670, Sigma-Aldrich) and 0.1% v/v Triton X-100 (108603, Sigma-Aldrich) in PBS. A volume of 10 µl medium supernatant was added to 90 µl p-nitrophenyl phosphate buffer and incubated for 90 min at 37°C. To stop the conversion of p-nitrophenyl phosphate into p-nitrophenol, 100 µl 0.3 M NaOH (106498, Sigma-Aldrich) was added. Absorbance was measured at 405 nm using a plate reader (Synergy^TM^ HTX, Biotek, Winooski, VT, USA). Standards with known p-nitrophenol concentrations, ranging between 0 and 0.9 µmol/ml, were used to calculate the TRAP activity per explant.

### 2.12 DNA quantification

DNA content was determined for the bone explants in dynamic (n=8) and static (n=6) conditions at the end of the 4-week culture period. Bone explants were cut in one half and two quarters. A quarter piece of the explant was two times washed with PBS to remove remaining medium, was weighted and placed into a tube containing 2 mL sterile water. Samples underwent 3 freeze-thaw cycles from -80°C to RT followed by 3 rounds of sonication (Soniprep 150, MSE, France) for 15 sec at 23 kHz with an amplitude of 12 µm and placement on ice between the cycles. After centrifugation at 3000 g for 10 min, DNA content was determined using Qubit^TM^ dsDNA HS Assay Kit (Invitrogen, USA). Briefly, 10 µl sample was added to 190 µl working solution prepared according to the manufacturer’s instructions and measured using the Qubit Fluorometer.

### 2.13 Measurement of alkaline phosphatase activity

To measure osteogenic activity, alkaline phosphatase activity (ALP) was quantified for bone explants in dynamic (n=8) and static (n=6) conditions. A quarter piece of the explant was washed two times with PBS to remove remaining medium, was weighted and placed into a tube containing 2 mL 0.2% v/v Triton X-100 (108603, Sigma-Aldrich) and 5 mM MgCl_2_ (M2393, Sigma-Aldrich). After incubation for 2 hours, samples were sonicated (Soniprep 150, MSE, France) three times for 15 sec at 23 kHz with an amplitude of 12 µm with placement on ice between the cycles. After centrifugation, ALP activity was quantified in supernatant by adding 100 μl substrate solution (10 mM p-nitrophenyl-phosphate (pNPP) (71768, Sigma-Aldrich), 0.15 M 2-amino-2-methyl-1-propanol (AMP) (A65182, Sigma-Aldrich) in UPW) to 20 μl of 0.75 M AMP and 80 μl sample. After 15 minutes at RT, the enzymatic reaction of pNPP into p-nitrophenol was stopped by adding 100 µl 0.2 M NaOH (106498, Sigma-Aldrich). Absorbance was measured at 450 nm at the start of the incubation period and after 15 minutes using a plate reader (Synergy^TM^ HTX, Biotek, Winooski, VT, USA). ALP activity per explant was calculated using standards with known concentrations of p-nitrophenol, ranging between 0 and 0.9 µmol/ml.

### 2.14 Statistical Analyses

Statistical analyses were performed, and graphs were prepared in GraphPad Prism (version 9.5.0, GraphPad, La Jolla, CA, USA). Data was tested for normal distribution using Shapiro-Wilk tests and homogeneity of variances was assessed from the residual plots. When these assumptions were met, a parametric test was performed. To compare between experimental groups on a specific timepoint a t-test or one-way ANOVA was performed, which was followed by Tukey’s post-hoc analysis to determine significant differences. If assumptions of normality and equal variances were not met, a non-parametric Kruskal-Wallis test followed by Dunn’s multiple comparisons test was performed to compare between experimental groups. To compare within a group over time, repeated measures ANOVA was performed, followed by Tukey’s post hoc test. Data is represented as mean ± standard deviation. A p-value of p<0.05 was considered statistically significant.

## 3. Results

### 3.1 Longitudinal µCT revealed bone formation mainly at the surface of osteochondral explants after static culture

The culture system for osteochondral explants allowed successful scanning of bone formation and resorption with weekly intervals. However, over the course of a week, changes in formation and resorption were nearly undetectable in static culture of osteochondral explants (**Figure S3A**). Larger time intervals needed to be taken between registered scans to make active bone remodeling apparent (**Figure S3B-C**). When an interval of 6 weeks was taken, new bone formation (6.48% ± 0.71) was predominantly observed at the outer surfaces of the bone (**Figure 2A**). Resorption was less abundant compared to formation (1.47% ± 0.21) and mainly seen as larger clusters at the edges of the sample. These clusters probably resulted from bone debris caused by the preparation procedure which were either lost during culture and consequently appeared as resorption clusters, or similar shaped clusters appeared in both resorption and formation channels due to small movement of the debris. Towards the center of the explant, formation and resorption speckles were observed in comparable amounts and evenly distributed throughout the bone. Much of this was identified as noise because colored voxels appeared inside trabeculae instead of on surfaces where bone remodeling was expected. Net bone remodeling showed a positive percentage of 5.01% ± 0.71 of volume gained during the 6-week culture period (**Figure 2B**). Histology confirmed µCT outcomes as collagen deposition was noticed at the outer surfaces of samples after 6 weeks culture (**Figure 2C-F**).

**Figure 2.**
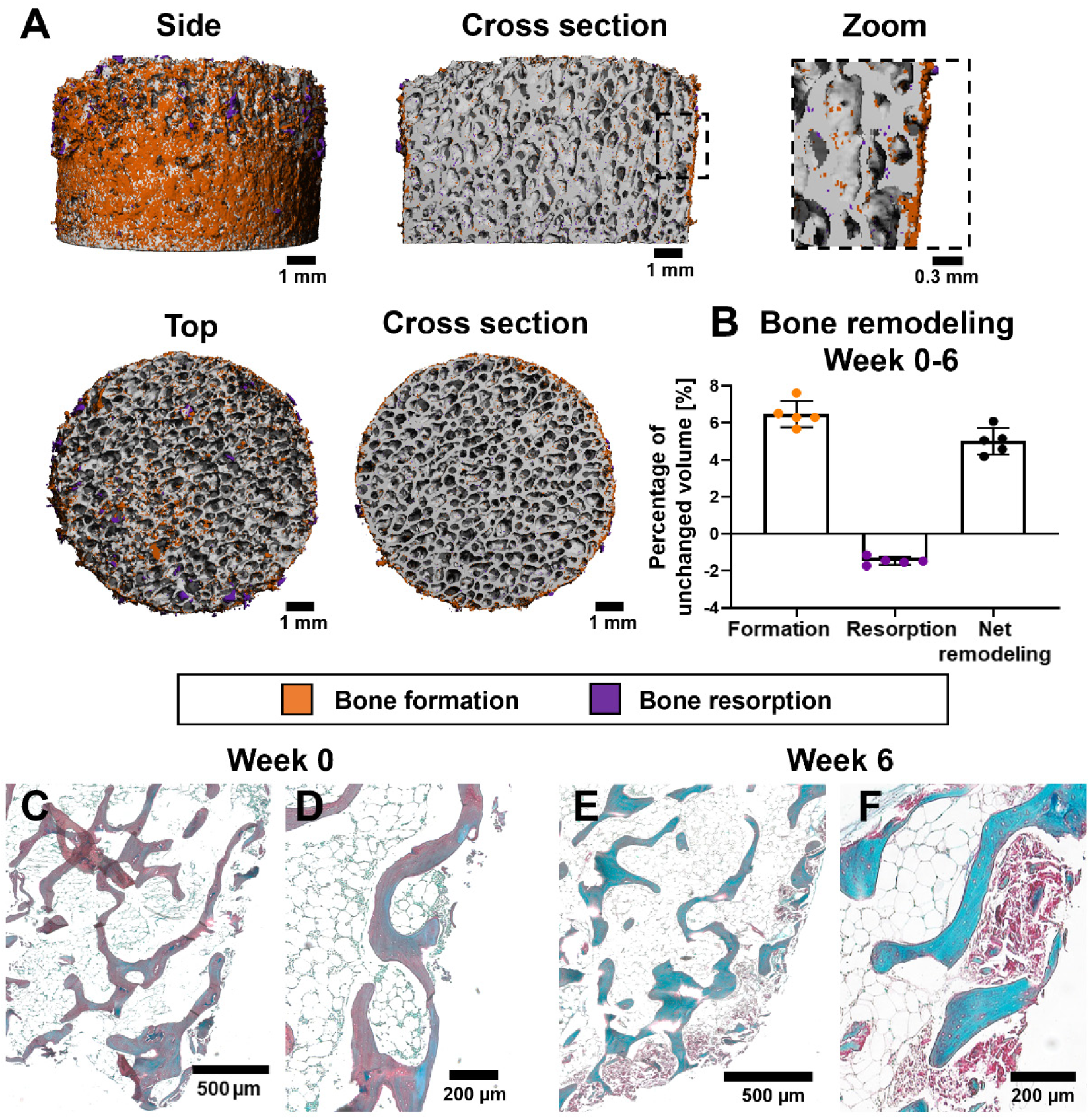
– Bone remodeling within osteochondral explants between day 0 and week 6 cultured in static condition. Bone formation (orange) and clusters of bone resorption (purple) at the outer surface of explants (A). Quantitative levels of bone formation and resorption as percentage of unchanged volume resulted in positive net remodeling after 6 weeks of culture (B). RGB trichrome stained sections visualized matrix deposition (collagen in red) at the outer surface which was absent at the start of culture (C, D) and developed during culture (E, F). Abbreviation: Picrosirius Red, Fast Green, Alcian Blue (RGB).

At the end of culture, osteochondral explants demonstrated to preserve trabecular architecture and cellularity in marrow spaces (**Figure 3A-B**). Remarkably, within the trabecular structures, white acellular tissue areas were observed within trabecular structures, with more pronounced occurrences closer to the subchondral bone (**Figure 3C**). Cells lining trabecular surfaces as well as osteocytic cell bodies in the lacunae were observed (**Figure 3B-C**). When specifically evaluating cells important for bone remodeling processes, MSC-like cells, positive for CD271 (**Figure 3D**), and osteoblast-like cells, positive for osteocalcin (**Figure 3E**), were identified in the explants. Osteoclasts were not observed at the end of culture. Measurement of TRAP activity confirmed the absence of active osteoclasts as TRAP levels decreased to similar levels as found in blank medium (0.2 µM pNPP/min) within 10 days of culture (**Figure 3F**).

**Figure 3.**
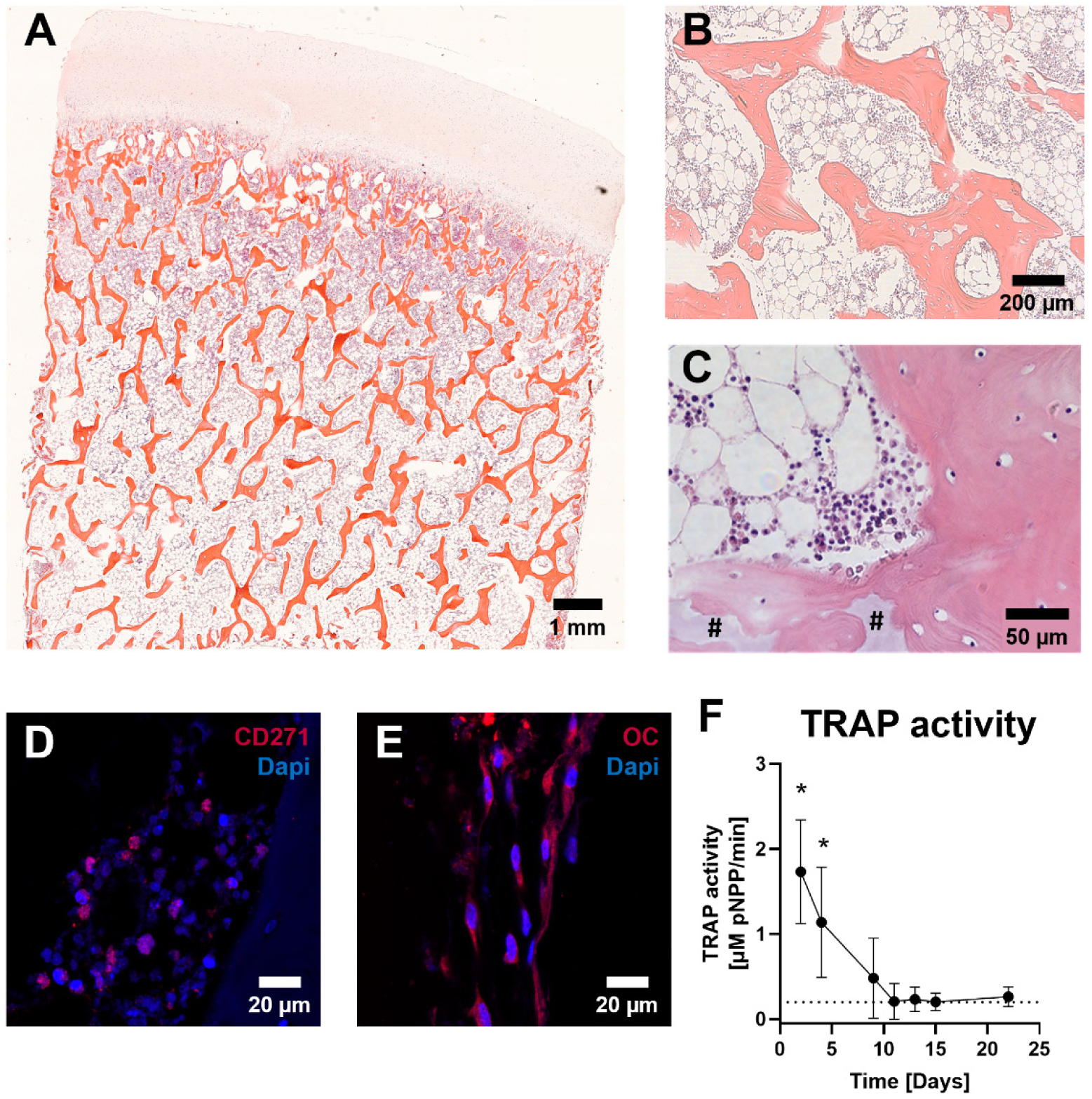
– Preservation of cells within osteochondral explants after 6 weeks of culture in static conditions. H&E-stained sections with well-preserved structural integrity and cellular morphology within the osteochondral tissue (A-C) including bone marrow cells between trabecular structures (B) and osteocytes within their lacunae (C). White acellular tissue observed within trabecular structures (#). Visualization of osteoprogenitor-like cells, identified with marker CD271 (D), and osteoblast-like cells, identified with marker osteocalcin (E). TRAP activity levels, as a measure of osteoclast activity, decreased towards levels of medium control (dotted line) within 10 days of culture (F). Data represents mean ± SD (*p<0.05 for timepoint day 2 and 4 with all following timepoints, except between day 2 and day 4 (not significant)). Abbreviations: hematoxylin and eosin (H&E), osteocalcin (OC) and tartrate-resistant acid phosphatase (TRAP).

### 3.2 No adverse effects of dynamic compressive loading on explant viability

To be able to calculate strain experienced by explants upon application of a certain force, apparent Young’s modulus of bone explants was determined. The moduli of trabecular porcine bone cores showed to vary between 16.7 and 142.8 MPa with a mean ± SD of 75.1 MPa ± 37.9 (**Figure 4A**). Statistically significant differences in Young’s moduli were found between donors. Application of cyclic compressive load of 40 N at 1 Hz using the bioreactor resulted in a displacement of the pistons which varied between 50 and 150 µm (**Figure 4B-C**). No adverse effects of this loading regime on DNA content, cell metabolic activity or cell death were observed (**Figure 4D-F**). Both statically and dynamically cultured bone explants showed significantly higher levels of cell death during the first 4 days of culture, which stabilized towards low levels from day 7 onwards (**Figure 4F**). Mean DNA content at the end of culture revealed more mean DNA for dynamically cultured explants compared to static culture, although this was not a statistically significant difference (**Figure 4D**).

**Figure 4.**
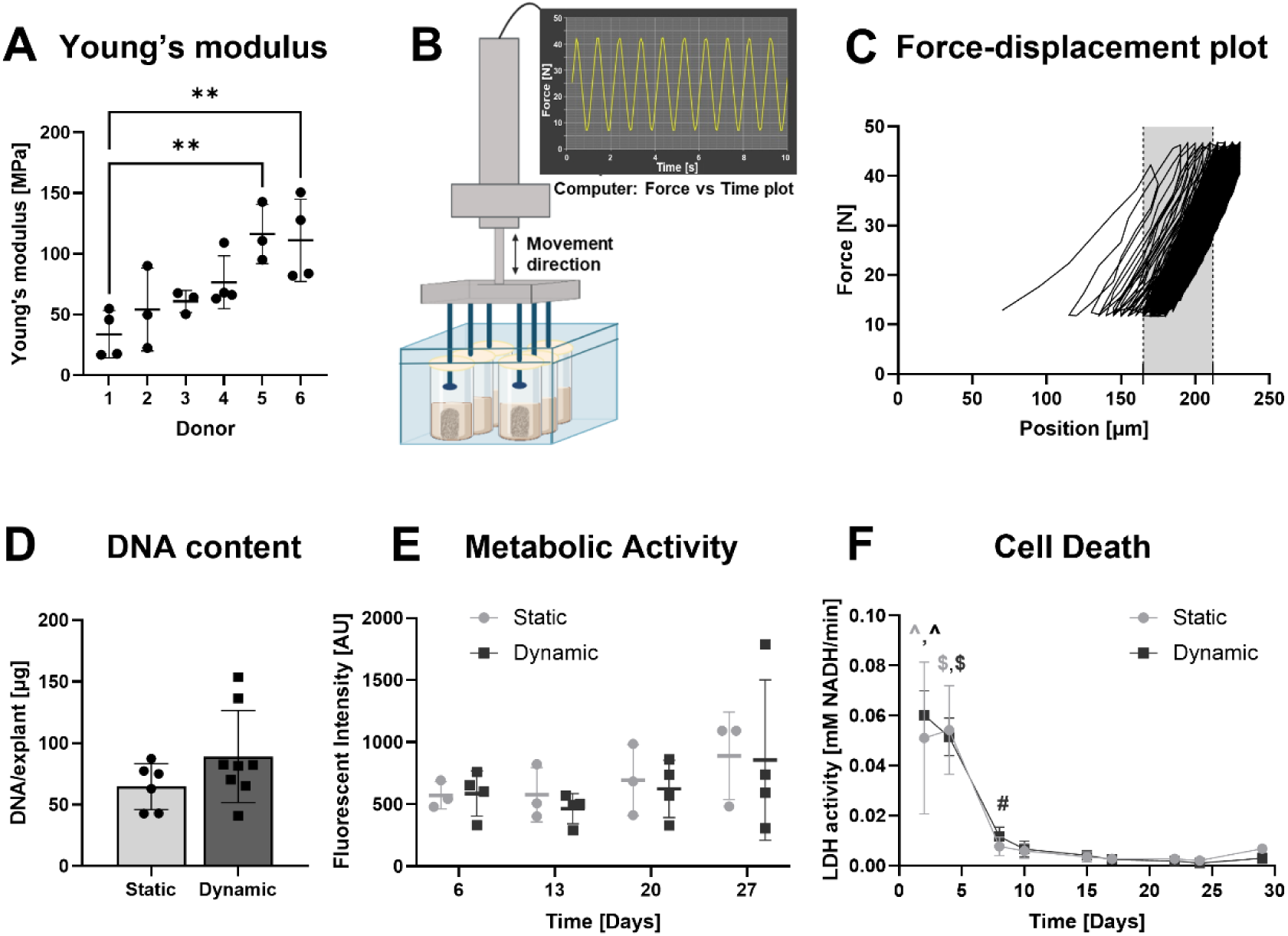
– Dynamic compressive loading applied to bone explants over a 4-week culture period. Young’s modulus of bone explants determined for 6 different porcine donors (A). Schematic of compression bioreactor with software generated force-time plot showing the first 10 seconds of the loading regime applying 40 N force at a frequency of 1 Hz for a total time of 30 minutes per day, 5 days a week, onto bone explants (B). Representative example of displacement of the pistons during compression cycles with grey area depicting displacement during 1 compression cycle, which was 50 µm in this case (C). No adverse effects of loading on DNA content measured for explants in static and dynamic conditions at the end of culture (D) as well as on metabolic activity (E) or cell death (F) monitored during culture were noticed. Data represents mean ± SD (**indicates p<0.01, ^ indicates p<0.0001 for timepoint day 2 with timepoints day 8, 10, 15, 17, 22, 24, 29, $ indicates p<0.0001 for timepoint day 4 with all other following timepoints, # indicates p<0.05 for timepoint day 8 with timepoints day 17, 22, 24 and 29). Abbreviation: lactate dehydrogenase (LDH).

### 3.3 Dynamic compressive loading altered bone remodeling responses

After registration of week 4 µCT images onto baseline images, it was observed that dynamically cultured samples showed increased amounts of formation in central areas of the bone explants (**Figure 5C-D**). In line with the osteochondral explants, statically cultured samples showed limited amounts of remodeling inside explants and bone formation was restricted to the outer surfaces for these samples (**Figure 5A**). Loaded samples demonstrated large resorption clusters at the outer shell (**Figure 5B**). The corresponding quantitative data revealed increased, but not statistically significant, amounts of bone resorption upon loading (static: 1.05% ± 0.2, loading: 2.60% ± 0.7) (**Figure 5E**). These large clusters of bone resorption at the outer surface in the dynamically loaded group were likely to be caused by scratching of the sides when taking the samples out of the bioreactor inserts. This did not happen in static condition as these explants were cultured in a regular well plate without inserts. Because this influenced the bone remodeling results, registration was repeated for the inner volumes only, excluding the outer 0.5 mm of the cores. A trend, not significant, towards an increased amount of bone formation was observed for explants cultured under dynamic loading (static: 1.13% ± 0.3, loading: 1.69% ± 0.38). Although differences were small, this trend was still visible in net remodeling (**Figure 5F**), where dynamically cultured samples gained more than five times higher volume during the culture period of 4 weeks compared to statically cultured explants (static: 0.08% ± 0.09, loading: 0.45% ± 0.25).

**Figure 5.**
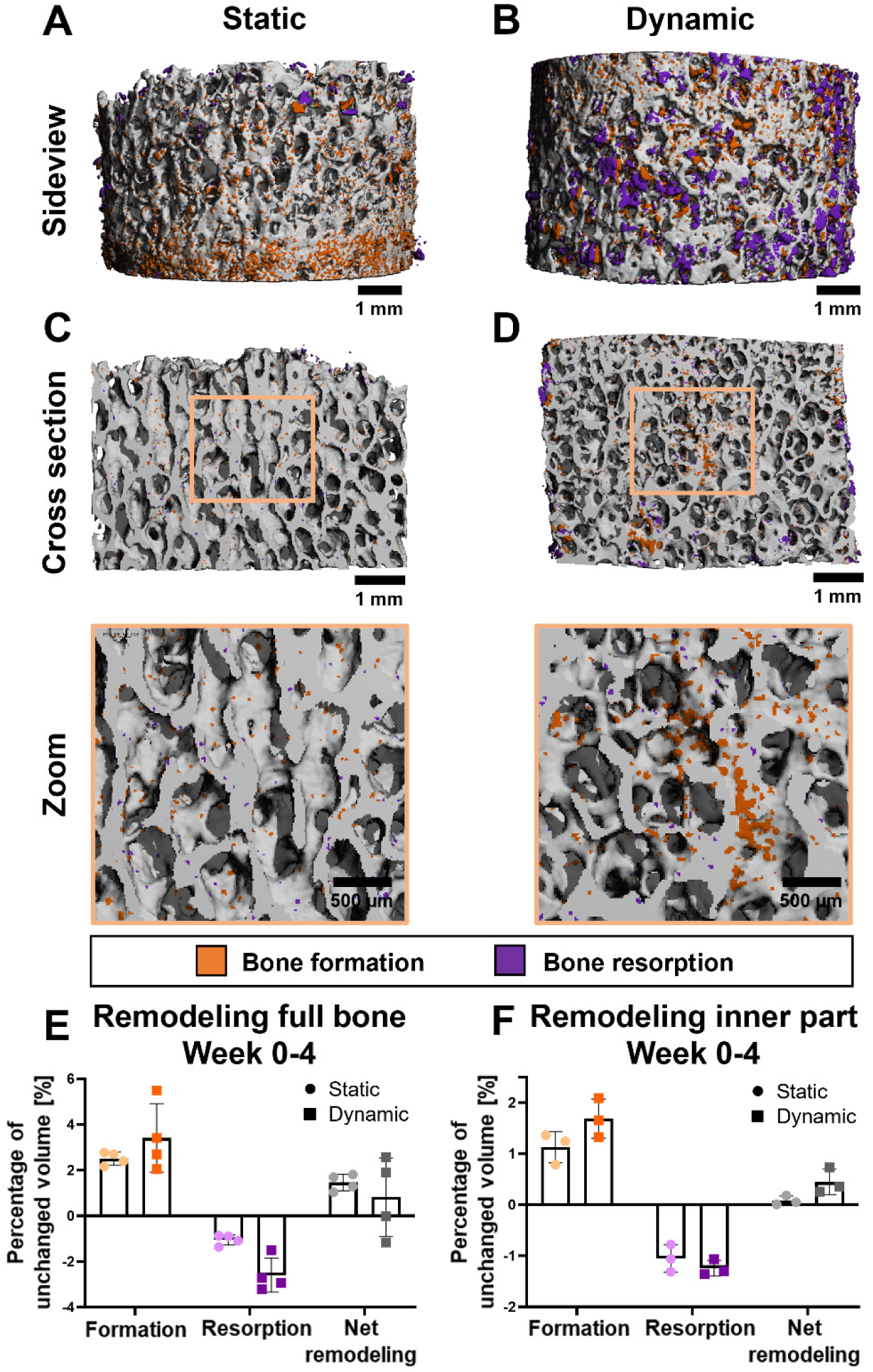
– Bone remodeling within bone explants over a 4-week culture period. Sideview of the outer surface of explants showing bone formation (orange) predominantly in static condition (A) and clusters of bone resorption (purple) in dynamic condition (B). Cross section of explant depicting enhanced bone remodeling within explants in dynamic (D) compared to static (C) culture conditions. Quantitative levels of bone formation and resorption as percentage of unchanged volume and net remodeling after registration of the complete bone (E) and for the inner part by excluding the outer 0.5 mm of the explant (F). Data represents mean ± SD.

Matrix deposition, in the form of collagen, in the central areas was only observed in loaded explants (**Figure 6A-B**). Mean ALP activity was higher for explants cultured in dynamic conditions compared to statically cultured samples, but this was not significantly different (static: 10.7 µM pNPP/min ± 4.3, loading: 14.7 µM pNPP/min ± 5.9) (**Figure 6C**). TRAP activity demonstrated that again osteoclasts did not maintain their activity during culture (**Figure 6D**).

**Figure 6.**
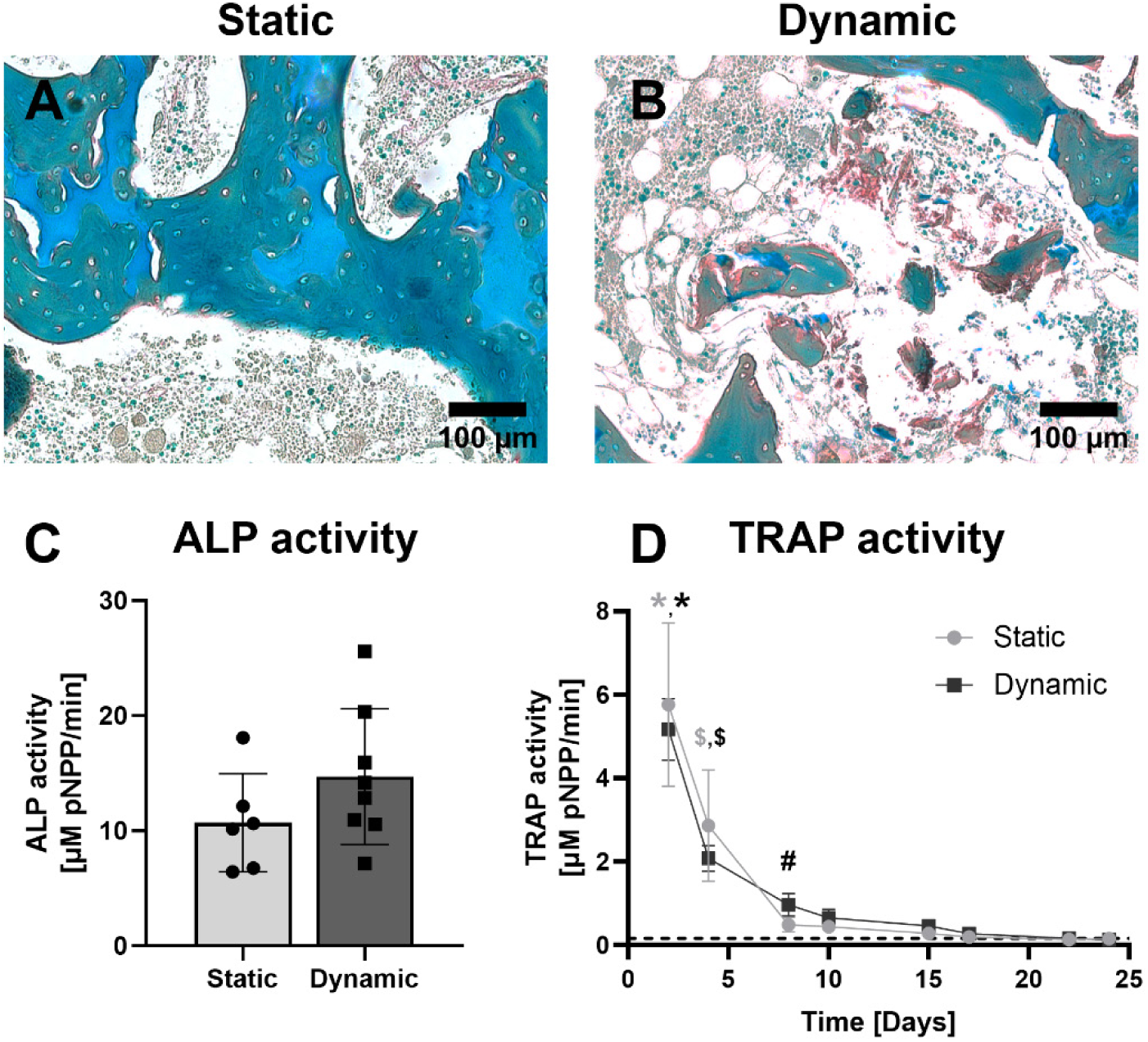
*-* Bone cell activity within bone explants cultured for 4 weeks. Matrix deposition (red) located at central parts of the explant, observed from RGB trichrome stained sections, was absent in static culture conditions (A) and present in dynamic culture conditions (B). No differences between static and dynamic culture conditions in osteoblast activity, measured by ALP activity in bone explants at the end of culture (C) and osteoclast activity, measured by TRAP activity in supernatant collected during culture (D). Data represents mean ± SD (* indicates p<0.0001 for timepoint day 2 with timepoints day 8, 10, 15, 17, 22, 24, 29, $ indicates p<0.001 for timepoint day 4 with all following timepoints, # indicates p<0.05 for timepoint day 8 with timepoints day 17, 22 and 24). Abbreviations: Picrosirius Red, Fast Green, Alcian Blue (RGB), alkaline phosphatase (ALP), tartrate resistant alkaline phosphatase (TRAP).

## 4. Discussion

With the preservation of tissue specific cells in their native 3D ECM, osteochondral explants have the ability to maintain direct communication between bone and cartilage thereby simulating an *in vivo* joint-like environment. So far, osteochondral explants are mainly utilized to study cartilage repair mechanisms and treatments, while there is a highly interconnected relationship with the underlying bone which is seen in for example osteoarthritis, which impacts both tissues. Therefore, it would be of interest to study remodeling processes in the bone part of osteochondral explants. Currently, bone remodeling is evaluated *ex vivo* with techniques including analysis of biochemical markers, histology and dynamic histomorphometry. Despite these techniques being valuable tools, they lack the spatiotemporal investigation of remodeling events, including both formation and resorption. Longitudinal µCT imaging and subsequent registration of consecutive scans represents an approach that has shown to be successful to evaluate spatiotemporal remodeling dynamics *in vitro* and *in vivo* (van Lenthe et al., 2007; Wehrle et al., 2019). To implement this method *ex vivo*, we developed a culture system for osteochondral explants that allowed for longitudinal monitoring of bone formation and resorption using µCT. It was hypothesized that implementation of this system, in which explants were cultured in a fixed position without the need to take out samples during culture, would facilitate successful registration of µCT images, thereby enabling the visualization and quantification of remodeling processes.

Effective registration without misalignment or registration errors was realized for the osteochondral explants within the presented culture system. Prior research in which bone explants were scanned pre- and post-experiment encountered issues in the registration using longitudinal µCT (Birmingham et al., 2015a). These issues were caused by motion artifacts and required manual transformation prior to registration, thereby limiting the volume that could be registered. The fixed position of the explants in our culture system achieved successful scanning and registration of the entire bone without limitations in volume. With weekly overlays it was demonstrated that remodeling could not be detected for the *ex vivo* cultures as weekly changes were very small and close to the resolution limit of µCT scanning. Larger intervals of 4-6 weeks between scans were needed to distinguish areas of active remodeling. In comparison, for *in vivo* studies, clear changes in bone remodeling within a mouse femur defect can be observed with weekly scan intervals (Wehrle et al., 2019). While the *in vivo* study had a higher spatial resolution of 10.5 µm compared to the *ex vivo* resolution of 16.4 µm, which could potentially reveal previously unseen remodeling sites, the lack of remodeling in the weekly scans could be explained by the slower dynamics of remodeling *ex vivo*. These slower dynamics are likely attributed to the absence of vascularization and blood flow in the *ex vivo* environment and should be considered in deciding on appropriate time-intervals between scans when working with explants.

The culture system demonstrated to be suitable for the *ex vivo* culture of osteochondral explants as evidenced by the preservation of viability, well-preserved trabecular architecture and cell morphology at the end of the culture period. Nonetheless, the elevated levels of cell death seen in the first days of culture indicated a decrease in cell viability. This observation seems inherent to the explantation of bone tissues as cells are exposed to hypoxia and mechanical stress during isolation procedures (de Vries-van Melle et al., 2012; Klüter et al., 2020; Stoddart et al., 2006). Thus far, it has not been investigated whether specific types of resident cells perish during explant isolation and if specific cell types remain viable. Regarding bone cells, this study revealed the absence of active osteoclasts but confirmed the presence of osteoblast-like cells, MSC-like cells and histologically intact osteocytes located within their lacunae at the end of the culture period. Incorporation of osteoclast activity, possibly through addition of fresh osteoclasts or monocytes, is an important point address in future research as it is a necessary element in studies of bone remodeling.

After 6 weeks in static culture, registration of endpoint µCT scans onto baseline scans revealed new mineralized volume predominantly at the outer surfaces of the explant. Histological analysis confirmed that this was indeed newly formed matrix as collagen was deposited in these regions. The activity restricted to outer surface is likely to be an effect of the static culture condition as nutrients in the medium do not easily reach the inner marrow spaces because of the diffusion limit and the hydrophobic nature of the bone marrow (Stoddart et al., 2006). This is in line with previous research that demonstrated a decrease in metabolic activity within the dense subchondral bone and in the central areas of osteochondral explants when cultured statically in the two-compartment system (Kleuskens et al., 2021; Schwab et al., 2017).

In the second part of this study, bone explants were subjected to compressive mechanical loading with the aim to capture increased bone formation. To ensure loading was experienced by the bone cells, the cartilage layer was removed. With only a bone core, the two compartments became unnecessary. Therefore, the culture chamber system was adapted to a one compartment system to be used for µCT scanning. We were able to successfully apply the compressive loading within the explant cultures, but levels of bone formation were not significantly increased in dynamically cultured samples compared to the statically cultured explants in our study. This was in contrast to previous *ex vivo* research where mechanical loading realized enhanced osteogenic responses within bone cores, observed from upregulated osteoblast activity, osteoid deposition, and trabecular thickening (Birmingham et al., 2015b, 2016; Chan et al., 2009; Curtis et al., 2018; David et al., 2008; Endres et al., 2009; Kogawa et al., 2018; Mann et al., 2006; Meyer et al., 2016; Simpson et al., 2009; Vivanco et al., 2013). A possible explanation for this might be that previous studies predominantly employed perfusion bioreactors along with compressive mechanical loading (David et al., 2008; Endres et al., 2009; Mann et al., 2006). In addition to the daily application of short periods of compression that were comparable to the culture setup used in our research, explants were continuously perfused during the culturing process. This may have provided continuous fluid flow as additional mechanical stimulus. Therefore, it is expected that a more pronounced effect on bone formation could be induced if perfusion is implemented in the culture system.

When remodeling was visually examined, areas of bone formation located towards the center of the explants were displayed upon addition of dynamic compressive loading, whereas bone formation remained restricted to outer edges within statically cultured explants. This might indicate that the compression-induced direct deformations and accompanying interstitial fluid flow in the explants were sensed by osteocytes locally, who started signaling towards osteogenic cells to induce bone formation. This would be in line with earlier work that showed that the application of mechanical loading did retain viability and function of osteocytes within cultured bone explants (Chan et al., 2009; Davidson et al., 2012). Still, the qualitatively observed gain in mineralized volume for dynamically cultured samples (in the center) was not reflected in the quantitative numbers of net remodeling (overall). This might be explained by the mounting of the explants in and out of the compression bioreactor inserts, thereby scratching the outer surface, causing large resorption clusters at the outer surface of explants, which did not appear for explants in static conditions as they were not cultured in inserts. By excluding the outer 0.5 mm for the registration for both loaded and non-loaded explants, it was possible to solely analyze remodeling processes in the inner core. Based on the absence of osteoclast activity it was assumed that the observed resorption was entirely noise in the inner part of the explants, and assuming that noise levels for formation and resorption would be similar, subtraction of resorption from formation corrected effectively for noise. The resulting parameter of net remodeling demonstrated a trend towards enhanced bone formation for explants subjected to compressive loading compared to static culture, where net remodeling was close to zero. More samples need to be included to verify the observed trend of increased bone formation upon compressive dynamic loading.

In this study it was shown that next to longitudinal µCT, a combination of evaluation methods was necessary to interpret remodeling results within the explants. Different analysis techniques, including histology, immunofluorescence, and biochemical characterization, were used to evaluate processes at a cellular level and thereby validated the longitudinal µCT results. In accordance with earlier studies using explants, osteogenic cells (osteoprogenitor-like cells, matrix synthesizing osteoblast-like cells, and osteocytes) were detected and could explain the levels found for bone formation (Davies et al., 2006; de Vries-van Melle et al., 2012; Mann et al., 2006). In contrast, osteoclast activity was not preserved in the explants, not even under dynamic culture conditions, indicating that the lost volume measured by µCT was not caused by active resorption of osteoclasts. The absence of osteoclast activity might be a result of the change from *in vivo* environment to an artificial *ex vivo* environment not being innervated and not having a vascular system, necessary to deliver new osteoclast precursors, which could evoke changes in signaling pathways potentially causing cell death. Furthermore, the culture medium did not contain osteoclast stimulating supplements. Factors such as RANKL, M-CSF, IL-6, TNF-α or bacterial factor LPS have been shown to stimulate osteoclastogenesis *ex vivo* (Cramer et al., 2021; Curtin et al., 2012; Sloan et al., 2013). Therefore, it could not be expected that osteoclast activity would have been preserved in the explants as the formation of osteoclast from their progenitors was not promoted. It would be of interest to evaluate and optimize components in the culture medium of explants and ideally realize culture medium conditions that do not favor any cell type such that cells could naturally interact with each other.

The compression bioreactor with the inserts was originally designed to create two compartments within the well-plate loaded in the bioreactor. In this study, the loading was exclusively used to promote osteogenesis in bone explants, making a second compartment redundant. Depending on the research question of future research, it could be of interest to use the compression bioreactor to apply compression onto osteochondral explants as it would mimic a joint-like environment more closely. Therefore, different loading regimes need to be tested and optimized because it would complicate loading with the load-bearing function of cartilage. Furthermore, cartilage is also subjected to sliding *in vivo* and would be valuable to test if such a sliding stimulus affects bone remodeling.

A limitation of the compression bioreactor used in this study was that it exerts compressive loads onto explants through a central computer-controlled stage connected to the single pistons, which results in the force being distributed over the explants. Consequently, the exact force that each explant received remains unknown, because it relies on the Young’s modulus, which varies between individual explants. Hypothetically, if Young’s moduli of the 4 explants that are simultaneously loaded would be similar, the force would be evenly distributed, and every explant would experience a force of 10 N with an imposed force of 40 N. For the average Young’s modulus measured for the porcine bone explants, 75.1 MPa, the application of a compressive force of 10 N resulted in strains > 2500 µstrain, which should stimulate bone formation according to the mechanostat theory (Frost, 2003; Kan et al., 2014). Previous research also demonstrated that bone explants revealed increased levels of osteoid deposition when subjected to strains > 2000 µstrains compared to non-loaded controls (Endres et al., 2009). However, if an explant receives a smaller force and/or has a lower Young’s modulus, the experienced strain could be lower than 2000 µstrain, potentially not resulting in the stimulation of bone formation. Overall, it would be of interest for future studies to explore different loading regimes and change to strain-controlled regimes as the bioreactor allows to provide a similar displacement to all explants. Furthermore, computer simulations could be used to model the load sensed by bone cells. For example, simulations of strain energy densities could be used to predict where bone formation and/or remodeling would be expected (Paul et al., 2018; Webster et al., 2015).

When working with bone tissue obtained from living specimens, donor variations should be considered. The Young’s moduli results emphasized the variation between porcine donors. Earlier findings reported Young’s moduli around 150 MPa for trabecular bovine bone cores of comparable volume, a stiffness that was approached by only a few samples in this study (David et al., 2008). Moreover, for human bone cylinders with comparable diameter, Young’s moduli of approximately 300 MPa for trabecular bone cores and approximately 15 GPa for the subchondral bone disks were reported (Li & Aspden, n.d., 1997). The relatively low Young’s moduli found in this study could be attributed to the young age of the pigs, which is 20 weeks ± 5 weeks. Pigs are only considered mature when they have an age of approximately 6 months. The acellular areas, that appeared white in histology, within the trabeculae might therefore be a result of bone tissue still undergoing endochondral ossification (Ferretti & Palumbo, 2021; Mackie et al., 2008). Furthermore, the variation in months might have led to the large range of Young’s moduli found. Young’s modulus values should be taken into account for further optimization of the bioreactor conditions, and it is recommended to determine the Young’s modulus for each sample before the start of the culture experiment. In future work, it would be advantageous to utilize a bioreactor system, such as the Zetos system, capable of measuring Young’s modulus, viscoelastic properties, fracture toughness, and fatigue properties of explants, in combination with finite element modeling to determine local stresses and strains (David et al., 2008; Davies et al., 2006; Endres et al., 2009).

For future studies, the presented *ex vivo* culture platform in combination with longitudinal µCT could be used to evaluate novel bone substitutes or drug treatments. This could aid in gaining increased knowledge of potential treatment effects. To enhance clinical relevance, it would be valuable to include human explant samples, which could be obtained from left-over bone tissue from surgeries, such as femoral heads or tibia plateaus (Swarup et al., 2018; Zankovic et al., 2021). Overall, the platform has the potential to be implemented in the preclinical testing phase of novel therapies preceding *in vivo* experimentation, thereby reducing and refining the number of animal studies needed.

## 5. Conclusion

This study presents a culture system platform suitable for the culture of osteochondral as well as bone explants and is simultaneously compatible with µCT scanning thereby enabling monitoring of remodeling processes. By registering consecutive scans, data on bone turnover is generated, which provides spatiotemporal information of remodeling dynamics. In conclusion, this culture system, when osteoclast activity could be implemented, has the potential to be a platform for standardized *ex vivo* evaluation of novel bone-related therapeutics, including materials for bone defects as well as bioactive agents for bone diseases.

## Author Contributions

E.C., K.H. K.I. and S.H. contributed to conception and design of the study and contributed to the methodology. L.K. provided expertise with the bioreactors. E.C. and K.H. performed experiments. E.C. analyzed results and wrote draft of the manuscript. E.C., K.H., L.K., S.H. and K.I. edited and reviewed the manuscript. All authors have read and approved the final submitted manuscript.

## Acknowledgement

The authors would like to thank Davy Wanders for the optimization of the bioreactor settings. The authors would also like to thank Jurgen Bulsink for manufacturing and maintenance of the culture chambers and tools for tissue isolation. This project received funding from ERC Proof of Concept (PoC) program BoneScreen with project number 956875 and research program TTW with project number TTW 016.Vidi.188.021, which is (partly) financed by the Dutch Organisation for Scientific Research (NWO). We gratefully acknowledge the Gravitation Program “Materials Driven Regeneration”, funded by the Netherlands Organization for Scientific Research (024.003.013).

## Conflict of interest statement

The authors declare that there are no conflict of interest.

## Notes

### Competing Interest Statement

The authors have declared no competing interest.

