## Supplementary information for "Culture system for longitudinal monitoring of bone remodeling *ex vivo*"

### Supplementary Figures

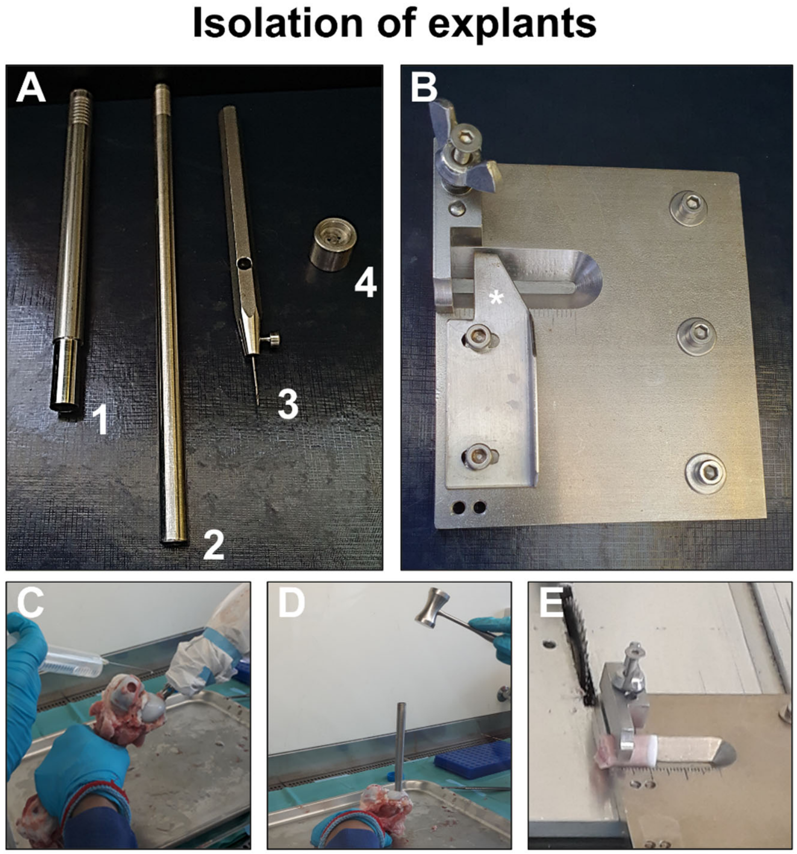


**Figure S1** – Equipment used for the isolation of explants. Custom-made tools (A) with hollow break-off tool (1), stick to remove explant from hollow break-off tool (2), Ø 1 mm drill (3) to create two holes within bone explants for mounting on the holder and guidance tool for drilling of the two holes (4). Custom-made stage (B) equipped with a scalebar and a removable component (*) used in combination with the bench saw to obtain explants with similar heights. The isolation process started with drilling, which was performed under constant cooling (C). After drilling, the break-off tool was hammered into the femoral condyles (D). After isolation of explants, the surface was cut straight (optionally: cartilage could be removed to obtain bone explants).

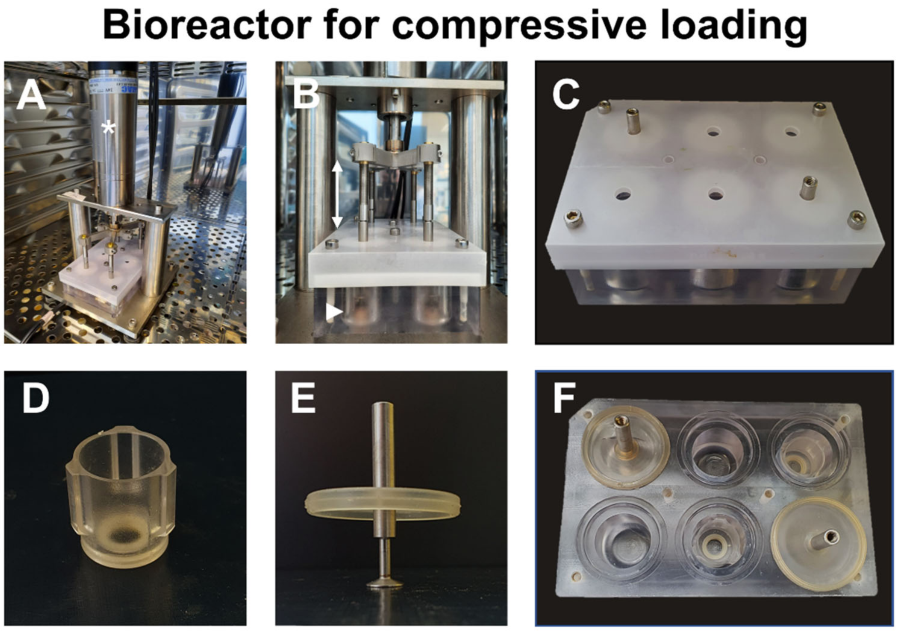


**Figure S2** – Compression bioreactor and accessory components. Bioreactor with linear actuator (*) within incubator to perform compression experiments at 37°C, 5% $CO_2$ (A). Linear actuator was connected to a central moving stage (moving in the direction of the double arrow) which was connected to 6 pistons to enable compression forces onto 6 explants (arrowhead indicates explant) at the same time (B). Custom-made well plate (C, F) in which inserts (D, F) were placed. Wells were sealed with silicon rubber where the piston penetrated trough (E, F). A hard plastic lid closed off the complete well plate (C).

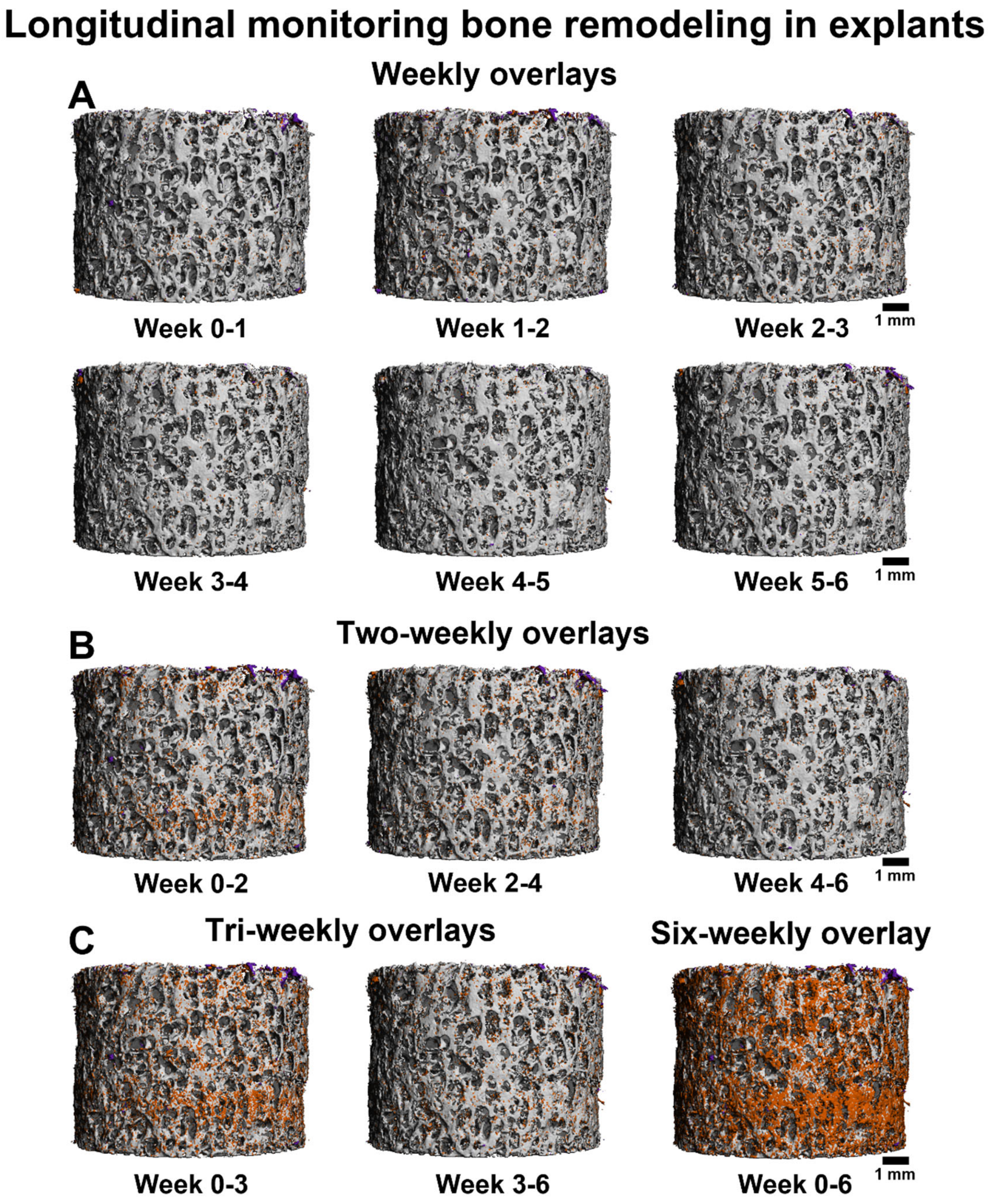


**Figure S3** – Representative example of bone remodeling in an osteochondral explant cultured for 6 weeks. The interval between registered scans was enlarged from weekly (A), two-weekly (B), tri-weekly (C) to six-weekly (D) to illustrate detectable bone remodeling with bone formation (orange), bone resorption (purple) and unchanged volume (grey).
